# Synaptic plasticity of inhibitory synapses onto medial olivocochlear efferent neurons

**DOI:** 10.1101/2022.01.05.475100

**Authors:** Lester Torres Cadenas, Hui Cheng, Catherine J.C. Weisz

## Abstract

The descending auditory system modulates the ascending system at every level. The final descending, or efferent stage, is comprised of lateral olivocochlear (LOC) and medial olivocochlear (MOC) neurons. MOC somata in the ventral brainstem project axons to the cochlea to synapse onto outer hair cells (OHC), inhibiting OHC-mediated cochlear amplification. MOC suppression of OHC function is implicated in cochlear gain control with changing sound intensity, detection of salient stimuli, attention, and protection against acoustic trauma. Thus, sound excites MOC neurons to provide negative feedback of the cochlea. Sound also inhibits MOC neurons via medial nucleus of the trapezoid body (MNTB) neurons. However, MNTB-MOC synapses exhibit short-term depression, suggesting reduced MNTB-MOC inhibition during sustained stimuli. Further, due to high rates of both baseline and sound-evoked activity in MNTB neurons *in vivo*, MNTB-MOC synapses may be tonically depressed. To probe this, we characterized short-term plasticity of MNTB-MOC synapses in mouse brain slices. We mimicked *in vivo*-like temperature and extracellular calcium conditions, and *in vivo*-like activity patterns of fast synaptic activation rates, sustained activation, and prior tonic activity. Synaptic depression was sensitive to extracellular calcium concentration and temperature. During rapid MNTB axon stimulation, post-synaptic currents (PSCs) in MOC neurons summated but with concurrent depression, resulting in smaller, sustained currents, suggesting tonic inhibition of MOC neurons during rapid circuit activity. Low levels of baseline MNTB activity did not significantly reduce responses to subsequent rapid activity that mimics sound stimulation, indicating that, *in vivo*, MNTB inhibition of MOC neurons persists despite tonic synaptic depression.

**Key points summary:**

- Inhibitory synapses from MNTB onto MOC neurons exhibit short-term plasticity that is sensitive to calcium and temperature, with enhanced synaptic depression occurring at higher calcium concentrations and at room temperature
- High rates of background synaptic activity that mimic the upper limits of spontaneous MNTB activity cause tonic synaptic depression of MNTB-MOC synapses that limits further synaptic inhibition
- High rates of activity at MNTB-MOC synapses cause synaptic summation concurrent depression to yield a response with an initial large amplitude that decays to a tonic inhibition

## Introduction

Mammalian auditory systems encode acoustic stimuli that occur over many orders of magnitude of sound intensity. Animals require mechanisms to shift the gain of the auditory system between softer and louder sounds to both enable soft sound detection and to prevent system saturation during loud sounds. The medial olivocochlear (MOC) neurons are one of multiple gain control systems that adjust auditory sensitivity (Galambos, 1956; Desmedt, 1962; Wiederhold and Peake, 1966; Wiederhold and Kiang, 1970; Geisler, 1974; Guinan and Gifford, 1988). MOC neuron somata are located in the ventral brainstem, and their axons project to the cochlea to form cholinergic synapses onto outer hair cells (OHC) (reviewed in Guinan, 1996). Through coupling of cholinergic responses in the OHC to SK or BK potassium channels, OHCs are inhibited by MOC activity (reviewed in Fuchs and Lauer, 2019). This inhibition suppresses the OHC “cochlear amplifier” effects, resulting in reduced OHC activity, and in turn, reduced auditory nerve activity with broadened tuning curves (Galambos, 1956; Fex, 1962a; Wiederhold and Kiang, 1970; Mountain, 1980; Siegel and Kim, 1982; Art et al., 1985). Thus, the MOC system is implicated in cochlear gain control, as well as detection of salient sounds, protection from noise trauma, and auditory attention (Oatman, 1976; Glenn and Oatman, 1977; Winslow and Sachs, 1987; Rajan, 1988, 1995; Kawase et al., 1993; Reiter and Liberman, 1995; Delano et al., 2007; Taranda et al., 2009; Tong et al., 2013; Maison et al., 2013; Terreros et al., 2016; Boero et al., 2018) via their actions on OHCs.

The patterns of activity of MOC synapses onto OHCs in the cochlea are governed by MOC somatic activity, which is in turn determined by interplay between intrinsic electrical neuronal properties and synaptic inputs that provide information about sound stimuli. *In vivo* recordings from MOC axons demonstrate low sound thresholds, narrow, V-shaped tuning curves qualitatively similar to auditory nerve tuning curves, and short-latency responses to loud sounds (Fex, 1962b; Robertson, 1984; Robertson and Gummer, 1985; Brown, 1989; Maison et al., 2003). Many MOC neurons have zero to low rates of action potentials *in vivo* in the absence of sound (Robertson and Gummer, 1985; Brown, 1989; Maison et al., 2003). The pre-synaptic circuitry driving MOC neuron activity is incompletely known, but anatomical (Brown et al., 2003, 2013; De Venecia et al., 2005; Darrow et al., 2012) and recent functional (Romero and Trussell, 2021) work indicates that T-stellate cells of the ventral cochlear nucleus (VCN) are the primary sound-driven inputs to MOC neurons, with contributions from cells of the cochlear nucleus (CN) small shell cap (Hockley et al., 2021). MOC neurons are also strongly excited by descending projections from the inferior colliculus (IC) (Romero and Trussell, 2021). In contrast to these excitatory synaptic inputs, we previously demonstrated that MOC neurons receive inhibitory synaptic inputs from neurons of the Medial Nucleus of the Trapezoid Body (MNTB), which can suppress spontaneous action potentials in MOC neurons in brain slice preparations *in vitro* (Torres Cadenas et al., 2020). MOC neurons may also receive additional synaptic inputs from brain regions involved in sound perception, as well as those involved in other roles such as attention (Faye-Lund, 1986; Caicedo and Herbert, 1993; Thompson and Thompson, 1993; Vetter et al., 1993; Mulders et al., 2002; Mulders and Robertson, 2002; Groff and Liberman, 2003; Horvath et al., 2003; Ota et al., 2004; Gómez-Nieto et al., 2008a; Brown et al., 2013; Christian Brown et al., 2013; Suthakar and Ryugo, 2016). How these various synaptic inputs integrate and impact MOC neuron activity remains an important unanswered question in auditory processing and cochlear gain control.

MNTB neurons are an integral component of auditory brainstem circuitry, forming inhibitory synapses onto not only MOC neurons in the ventral nucleus of the trapezoid body (VNTB), but also onto other neurons of the superior olivary complex (SOC) including lateral superior olive (LSO), medial superior olive (MSO), nuclei of the lateral lemniscus (NLL) and the superior peri-olivary nucleus (SPON) (Glendenning et al., 1981; Kuwabara and Zook, 1991, 1992; Banks and Smith, 1992; Smith et al., 1998). MNTB neurons are strongly activated by VCN Globular Bushy Cells (GBC) which synapse onto the MNTB somata with the large calyx of Held axon terminal (Held, 1893; Ramón y Cajal, 1909; Friauf and Ostwald, 1988; Spirou et al., 1990; Kuwabara et al., 1991; Smith et al., 1991) to form a well-characterized high-fidelity synapse (Guinan and Li, 1990; Wu and Kelly, 1993; Barnes-Davies and Forsythe, 1995; Borst et al., 1995; Taschenberger and von Gersdorff, 2000) but see (Kopp-Scheinpflug et al., 2003). *In vivo*, MNTB neurons have baseline spiking activity in the absence of sound that has been measured from 0.15 to 190 Hz, with averages around 10-20 Hz (Sommer et al., 1993; Smith et al., 1998; Kopp-Scheinpflug et al., 2003; Hermann et al., 2007; Kadner and Berrebi, 2008). MNTB neuron responses to sound onset exhibit a “primary-like with notch” pattern peri-stimulus time histogram (PSTH) (Guinan and Li, 1990; Smith et al., 1998; Paolini et al., 2001; Kopp-Scheinpflug et al., 2008; Tolnai et al., 2009). MNTB neuron sound-evoked spike rates can reach to the hundreds of Hz (Kopp-Scheinpflug et al., 2008). How this fast spike rate of MNTB neurons inhibits MOC neurons to shape auditory processing is unclear.

To increase our understanding of how MOC neurons adjust their activity over the course of auditory stimuli, we characterized the short-term plasticity of the MNTB-MOC synapse. We show that short-term plasticity is affected by temperature, specifically that synaptic depression is reduced at physiological temperature. We also demonstrate that extracellular calcium concentrations that that are known to modulate pre-synaptic neurotransmitter release affect plasticity at this synapse. MNTB-MOC synapses exhibit depression at both physiological and high extracellular calcium concentrations, and exhibit facilitation at low extracellular calcium concentrations. By applying low rates of stimulation of MNTB axons to mimic baseline spontaneous *in vivo* activity we examine the subsequent MOC synaptic responses to higher rates of activity, such as might occur at sound onset. Stimulation of MNTB-MOC synapses at typical baseline spontaneous rates does not appreciably alter subsequent synaptic activity. Importantly, we show that stimulation of the MNTB-MOC synapse at higher rates of activity significantly reduces the subsequent effect of MNTB-MOC synapses due to synaptic depression. Finally, we test the response of MNTB-MOC synapses to long, fast stimulation to maximally depress the synapses. From this we demonstrate that MNTB neurons can maintain sustained inhibition of MOC neurons even during highly active periods. This sustained inhibition ultimately summates to reach a steady state to generate a tonic inhibition of MOC neurons.

## Methods

### Animals and Slice Preparation

Animal procedures followed National Institutes of Health guidelines, as approved by the National Institute of Neurological Disorders and Stroke / National Institute on Deafness and Other Communication Disorders Animal Care and Use Committee. Brain slices were prepared from post-natal day (P) 14-23 mice of either sex resulting from a cross between ChAT-IRES-Cre transgenic mice on a C57BL/6J background strain (Jackson Labs 028861), with tdTomato reporter mice (Ai14, Jackson Labs 007914). Hemizygotes were used to prevent deleterious effects noted in ChAT-IRES-Cre homozygotes (Chen et al., 2018). As noted in the mouse line descriptions, this ChAT-IRES-Cre strain is occasionally prone to ectopic expression of Cre in vasculature, glia, or non-cholinergic neurons due to the presence of the neo cassette, causing tdTomato labeling in non-cholinergic cells. If any ectopic expression patterns were observed, the tissue from the animal was not used. Mice were euthanized by carbon dioxide inhalation at a rate of 20% of chamber volume per minute, then decapitated. The brain was removed in cold artificial cerebrospinal fluid (aCSF) containing (in mM): 124 NaCl, 1.2 CaCl_2_, 1.3 MgSO_4_, 5 KCl, 26 NaHCO_3_, 1.25 KH_2_PO_4_, 10 dextrose. 1 mM kynurenic acid was included during slice preparation. The pH was equal to 7.4 when bubbled with 95% O_2_ / 5% CO_2_. 300-micron coronal brain slices containing nuclei of the SOC including the MNTB and VNTB were cut with a vibratome (Leica VT1200S) in cold aCSF. The slices were stored in a custom interface chamber at 32°C for one hour and then allowed to cool to room temperature for electrophysiological recordings. The slices were used within four hours of preparation.

### Patch-clamp electrophysiological recordings

Brain slices were transferred to a recording chamber continuously perfused at a rate of ∼2-3 ml/min with aCSF bubbled with 95% O_2_ / 5% CO_2_. The majority of voltage-clamp recordings were performed at physiological temperature (35 ± 1°C), although some were performed at room temperature (24 ± 1°C), where indicated. In experiments to test the effect of temperature on synaptic properties, the different temperatures were applied in random order. The slices were viewed using a Nikon FN-1 microscope with DIC optics and a Nikon NIR Apo 40X/ 0.80 N/A water-immersion objective. The images were collected with a QICLICK, MONO 12BIT camera (Nikon) using NIS-Elements software (Nikon). MOC neurons were identified for whole-cell voltage-clamp recordings by their position in the VNTB (MOC neurons in the dorsal peri-olivary region were not recorded from in this study) and visibility using red epifluorescence (546 nm emission filter, Lumencor Sola lamp). The recordings were performed using a MultiClamp 700B and DigiData 1440A controlled by Clampex 10.6 software (Molecular Devices). The recordings were sampled at 50 kHz and filtered on-line at 10 kHz. The internal solution for MOC recordings contained (in mM): 56 CsCl, 44 CsOH, 49 D-gluconic acid, 1 MgCl_2_, 0.1 CaCl_2_, 10 HEPES, 1 EGTA, 0.3-Na-GTP, 2 Mg-ATP, 3 Na_2_-phosphocreatine, 5 QX-314, 0.25% biocytin, 0.01 AlexaFluor-488 hydrazide. The Cl^-^ equilibrium potential was ∼-20 mV.

Recording pipettes were pulled from 1.5 mm outer diameter borosilicate glass (Sutter Instruments) to tip resistances of 3-6 MΩ. Series resistances were corrected 50-85%. The cells were voltage-clamped at −60 mV unless stated otherwise. Membrane voltages were not adjusted for a measured liquid junction potential of −2 mV (Cs gluconate solution). MNTB axons were electrically stimulated to evoke neurotransmitter release and generate post-synaptic currents (PSC) in MOC neurons by a large diameter glass pipette of ∼10 or ∼30 µm diameter filled with aCSF connected to an Iso-Flex Stimulus Isolation Unit (A.M.P.I.), placed in the MNTB axon bundle within or at the lateral edge of the MNTB. The stimulus intensity was increased until a maximal evoked PSC (ePSC) amplitude was reached, then was turned down to an intermediate intensity ∼200 µA. CNQX (5 µM) was included in all experiments to block excitatory transmission through AMPA-type glutamate receptors by addition to the recirculating aCSF solution. Drugs and chemicals were obtained from Fisher Scientific or Sigma Aldrich.

### Experimental Design and Statistical Analysis

Evoked PSC amplitudes were measured in Clampfit 10.6 (Molecular Devices) by manually placing cursors at the immediate onset of the PSC (for trains of stimuli, during the decay of the preceding PSC immediately prior to the stimulus artifact), and at the peak of the PSC. The latency to the PSC response was measured from the beginning of the stimulus artifact to the onset of the response. The PSC time constant of decay was measured from the peak of the evoked PSC to the baseline plateau and was fit with a single exponential. In some experiments to measure the effect of spontaneous background activity on synaptic plasticity of inhibitory synaptic inputs, MNTB axons were first stimulated for 20 pulses at 10 or 100 Hz, followed by stimulation at higher rates of 50 to 500 Hz. The amplitude of ePSCs, probability of recording an evoked PSC, and facilitation index (S1-Sx)/S1 was computed during synaptic responses to stimulus trains as in our previous work (Torres Cadenas et al., 2020).

All statistical analyses were performed using Origin v2020 (Origin Laboratories) or R-version-4.0.2 using packages “lme4” and “lmerTest”. Linear mixed models were fit by REML (restricted (residual) maximum likelihood). T-tests used Satterthwaite’s method [‘lmerModLmerTest’]. Data were examined for normal distribution using the Shapiro-Wilk test. Nonparametric tests were applied given the non-normal distribution of many variables studied. The differences between two independent groups were assessed using a Mann-Whitney U test. Differences between multiple groups were tested with a Wilcoxon Rank Sum test. To evaluate the effects of stimulation rates on short-term plasticity at MNTB-MOC synapses, linear mixed-effects models were run with facilitation index as the dependent variable, stimulation rates and trains of electrical stimulation as fixed-effect independent variables. The random effects include random intercept for neurons and random slope for stimulation rate. We used the Akaike information criterion (AIC) to assist with selecting the appropriate statistical model. A lower AIC value indicates better quality of fit. The differences were considered significant if p < 0.05 and indicated by asterisks in figures. For figures, single traces were lowpass filtered at 2 kHz and in cases where more than one voltage-clamp trace was averaged, the data were not filtered. Summary box plots indicate the median and quartiles, with 10 and 90 percentiles indicated by error bars. A square within the box plot indicates the mean. Individual data points are overlaid. All data are presented as median ± median absolute deviation unless otherwise stated. The figures were prepared in Origin v2020 and Adobe Illustrator CC 2021 (Adobe).

## Results

Whole-cell patch-clamp recordings were performed from identified MOC neurons in brain slices from ChAT-IRES-Cre x tdTomato mice by targeting red fluorescent neurons in the ventral nucleus of the trapezoid body (VNTB). Using this approach, post-synaptic currents (PSCs) that were evoked (ePSCs) by electrical stimulation of pre-synaptic MNTB axons (Figure 1A) occurred in MOC neurons that were consistent with previous recordings (Torres Cadenas et al., 2020). With the high chloride concentration used here (60 mM), both excitatory and inhibitory PSCs were inward at a holding potential of −60 mV. AMPA receptors were blocked in all experiments with CNQX (5 µM) to isolate inhibitory PSCs. After establishing this baseline, we tested whether there was a difference in strength of inhibitory synaptic inputs in male vs. female mice. We compared the amplitude of PSCs evoked by minimal stimulation (interpreted as the stimulation of a single axon) and found no difference (ePSC amplitude: female: 24.4 ± 7.25 pA, male: 28.1 ± 8.2, Mann-Whitney test U = 302, Z = −0.76, p = 0.44; ePSC time constant of decay: female: 11.6 ms ± 5.1 ms, male: 12.1 ± 4.5 ms, Mann-Whitney test U = 348, Z = 0.04487, p = 0.96; n = 23 female, n = 30 male). Due to a lack of difference in evoked PSCs of male vs female mice, mice of both sexes were pooled for further analyses.

**Figure 1.**
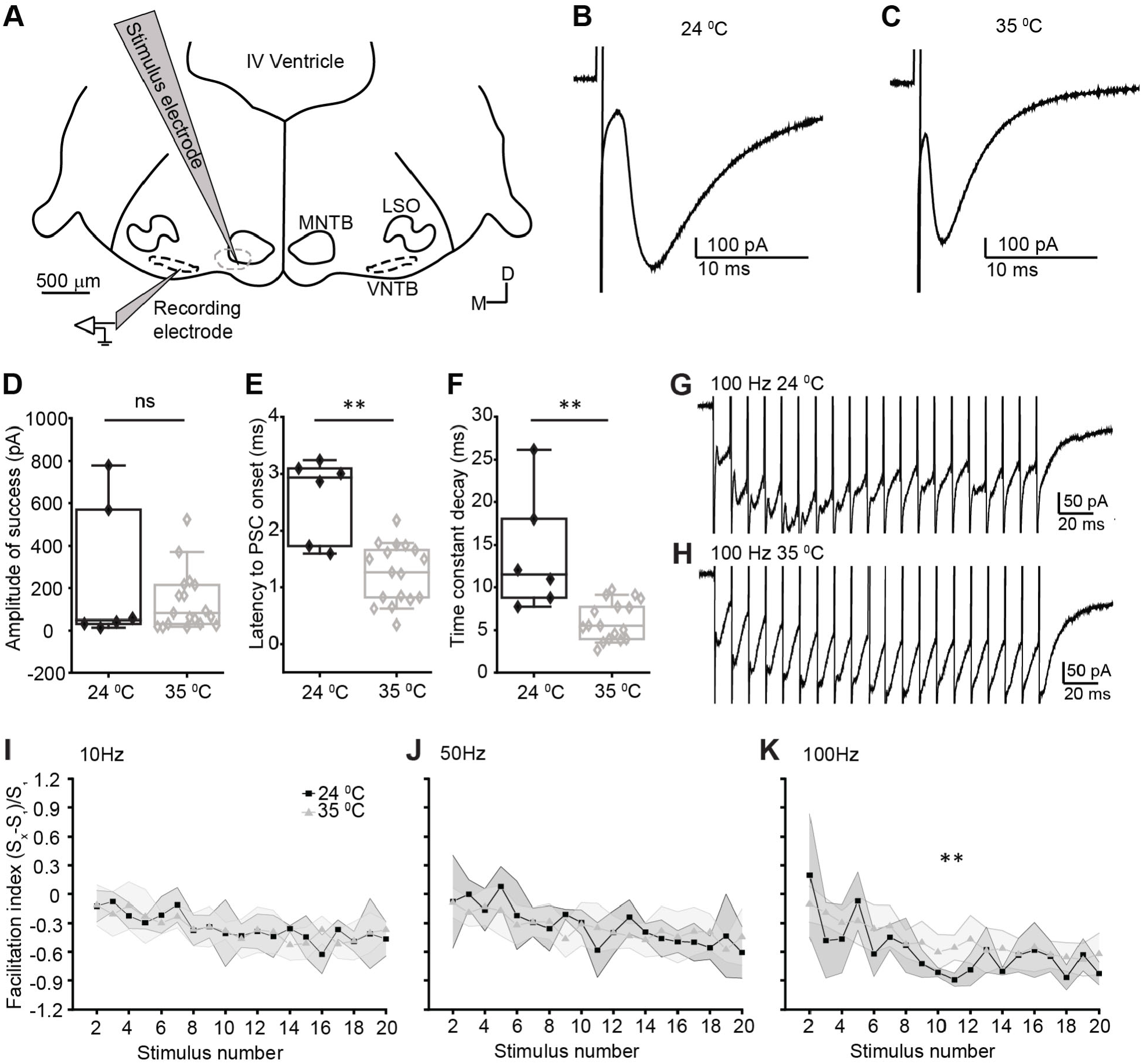
Temperature dependence MNTB-MOC synaptic responses. A. Schematic of experiments to characterize MNTB-MOC circuitry. B-C B. PSCs recorded in identified MOC neurons evoked by electrical stimulation of MNTB axons at A. RT (24 C) and B. PhT (35 C) D. Plot of PSC amplitude in RT and PhT, each dot represents the average from a single cell. E. Plot of latency from electrical stimulation of MNTB axons to evoked PSC onset in MOC neurons in RT and PhT. F. Plot of time constant of decay of PSCs evoked by electrical stimulation of MNTB axons in RT and PhT. G. Example traces of PSCs evoked by 100 Hz stimulation of MNTB axons at RT. H. Example traces of PSCs evoked by 100 Hz stimulation of MNTB axons at PhT. I-K. Facilitation index by stimulation number of PSCs evoked by stimulation of MNTB axons in RT vs PhT at I. 10Hz, J. 50 Hz, and K. 100 Hz. Black indicates RT, grey indicates PhT. Shading indicates MAD.

### Influence of temperature on synaptic plasticity

Previous MOC efferent neuron recordings from our group to characterize MNTB-MOC synapses were performed at room temperature (RT: ∼24 C) (Torres Cadenas et al., 2020). Because temperature can have diverse effects on properties of pre-synaptic vesicle release, post-synaptic responses, and synaptic plasticity, here we compared the effect of a higher, physiological temperature (PhT: ∼35 C), on post-synaptic responses in MOC neurons. During recordings from identified MOC neurons, MNTB axons were stimulated to release neurotransmitter, and ePSCs were measured in the MOC neuron. The amplitude of single ePSCs did not change between RT and PhT (RT amplitude: 48.75 ± 25.94 pA, n = 60 ePSCs in 6 neurons, PhT: 81.02 ± 62.35 pA, n = 170 ePSCs in 17 neurons, Mann Whitney U test, U = 49, Z = −0.11, p = 0.92, Figure 1A-D). However, after MNTB axon stimulation, MOC neurons had significantly shorter latencies to ePSC onset (RT ePSC latency: 2.93 ± 0.24 ms, n = 6 neurons, PhT: 1.27 ± 0.45 ms, n = 17 neurons, Mann Whitney U test, U = 92, Z = 2.84, p = 0.0045, Figure 2E) and had significantly faster time constants of decay in PhT compared to RT (RT ePSC tau: 11.51 ± 3.27 ms, n = 6 neurons, PhT: 5.53 ± 1.72 ms, n = 17 neurons, Mann Whitney U test, U = 95, Z = 3.05, p = 0.0023, Figure 2F). We next tested the short-term plasticity of MNTB-MOC ePSCs during trains of stimulation of MNTB axons at inter-stimulus intervals (ISIs) from 10 to 100 Hz at RT vs PhT (Figure 1G, H). We used the facilitation index to assess short-term synaptic plasticity. Using this index, a positive value indicates synaptic facilitation and a negative value indicates synaptic depression. We found that the facilitation index was not different across trains of 20 pulses between RT and PhT during 10 or 50 Hz stimulation (10 Hz RT: −0.44 ± 0.11, n = 6 neurons, 10 Hz PhT: −0.32 ± 0.09, n = 13 neurons, Mann-Whitney U Test: U = 136, Z = −1.28, p = 0.20; 50 Hz RT: −0.36 ± 0.08, n = 6 neurons, 50 Hz PhT: −0.34 ± 0.08, n = 13 neurons, Mann-Whitney U Test: U = 137, Z = −1.26, p = 0.21, Figure 2I, J). However, at 100 Hz, ePSCs depressed more at RT compared to PhT (100 Hz RT: −0.61 ± 0.13, n = 6 neurons, 100 Hz PhT: −0.44 ± 0.07, n = 13 neurons, Mann-Whitney U Test: U = 56, z = −3.62, p = 2.9E-4, Figure 2K). These results indicate that synaptic activity at MNTB-MOC synapses is more resistant to short-term depression at physiological temperatures.

**Figure 2.**
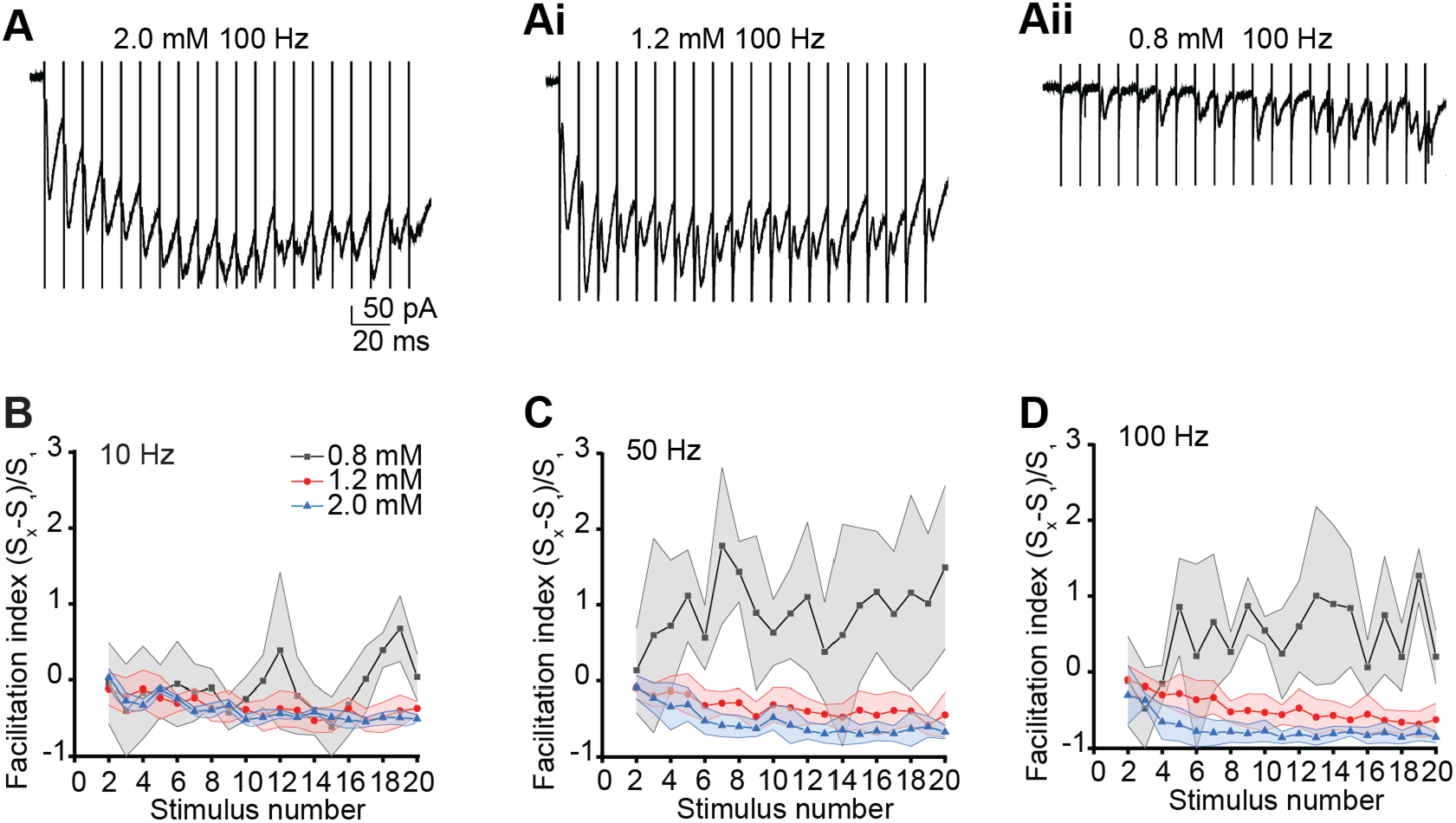
Calcium dependence of short-term plasticity at MNTB-MOC synapses. A. ePSCs evoked by trains of electrical stimulation of MNTB axons recorded in high (2.0 mM, A), physiological (1.2 mM, Ai) and low (0.8 mM, Aii) extracellular calcium. B-D. Facilitation index of PSCs evoked from MNTB axons in 0.8 mM (black), 1.2 mM (red) or 2.0 mM (blue) extracellular calcium, at B. 10Hz, C. 50 Hz, and D. 100 Hz stimulation rates. Shading indicates MAD.

### Calcium dependence of plasticity at MNTB-MOC synapses

Many auditory neurons in the CN and SOC fire rapid action potentials through the duration of a sound stimulus, so short-term synaptic plasticity due to high rates of synaptic activity has important implications for auditory neuron circuit function. Short-term synaptic plasticity is governed by mechanisms that affect both pre- and post-synaptic elements, including extracellular calcium concentrations (Neher and Sakaba, 2008; Regehr, 2012). To characterize calcium-dependence of short-term synaptic plasticity at the MNTB-MOC synapses, we applied trains of electrical stimulation to MNTB axons at rates of 10-100 ms in extracellular calcium concentrations from 0.5 to 2.0 mM calcium. The effect of stimulation rates was analyzed with a linear mixed-effect model. By including random intercept for neurons and random slope for stimulations rate, we assume that the facilitation index and the effect of stimulation rates vary for each neuron. This was the best fitting model (AIC = −456.95) compared to a model without random effects (AIC = −87.43) and a model with only a random intercept (AIC = −279.62). Similar to our previous work (Torres Cadenas et al., 2020), electrical stimulation-evoked PSCs exhibited synaptic depression in 2.0 mM extracellular calcium at stimulation rates of 10, 50, and 100 Hz, with 100 Hz stimulation evoking a significantly greater degree of depression relative to 10 Hz stimulation (*ß* = −0.35, S.E. : 0.086, *t* = −4.05, *p* = 0.0098, Figure 2A). Recent work suggests that an extracellular calcium concentration of 1.2-1.5 mM is thought to approximate *in vivo* levels (Borst, 2010). In 1.2 mM extracellular calcium, trains of stimulation of MNTB axons did not evoke synaptic depression at the second pulse in the train, consistent with previous work using paired pulse stimulation and an ISI of 10 ms (Torres Cadenas et al., 2020). However, with sustained synaptic activity through the train, synaptic depression was observed at stimulation rates of 10-100 Hz (One sample Wilcoxon signed-rank test indicated facilitation index was significantly less than zero, median = −0.398, V = 43775, *p* < 2.2e-16, Figure 2Ai). Synaptic depression was significantly greater in 2.0 mM compared to 1.2 mM extracellular calcium at train rates of 100 and 50, but not 10 Hz, as measured by facilitation index (100 Hz: 2.0 mM −0.299 ± 0.38, 1.2 mM −0.105 ± 0.18, Wilcoxon Rank Sum Test W = 29680, p = 2.2e-16; 50 Hz: 2.0 mM 0.084 ± 0.15, 1.2 mM 0.090 ± 0.15, Wilcoxon Rank Sum Test W = 18780, p = 3.4e-07; 10 Hz: 2.0 mM 0.029 ± 0.11, 1.2 mM −0.114 ± 0.21, Wilcoxon Rank Sum Test W = 19690, p = 0.27, n = 6 neurons, Figure 2B-D). These results indicate that short-term synaptic depression of MNTB-MOC synapses is not as strong in physiological conditions (1.2 mM calcium) compared to higher calcium (2.0 mM) conditions, but synaptic depression in 1.2 mM calcium is significant with sustained synaptic activity.

We further tested the calcium-dependence of short-term synaptic plasticity at MNTB-MOC synapses by lowering extracellular calcium concentrations to 0.5 or 0.8 mM. Evoked PSCs were rarely recorded in 0.5 mM calcium (1 ePSC in 5 trains of stimulations each in 6 neurons, not shown), so synaptic plasticity was not further addressed. However, the rate of PSCs evoked in 0.8 mM extracellular calcium was sufficient to assess plasticity. ePSCs occurred at the first pulse in a train with a lower probability in low (0.8 mM) extracellular calcium compared to 1.2 or 2.0 mM calcium (0.8 mM probability = 0.2 ± 0.2, n = 8 neurons, 1.2 mM probability = 1 ± 0, n = 17 neurons, 2.0 mM probability = 1 ± 0, n = 6 neurons; 0.8 mM vs 1.2 mM: Mann-Whitney U test: U = 0, Z = −4.34, p = 1.43E-5; 0.8 mM vs 2.0 mM: Mann-Whitney test: U = 0, Z = −3.12, p = 0.0018). When trains of electrical stimulation were applied to MNTB axons (Figure 2Aii), MNTB-MOC synapses facilitated at stimulation rates of 100 and 50 Hz, but not at 10 Hz (One sample Wilcoxon signed-rank test indicated the facilitation index was significantly greater than zero for 100 and 50 Hz: 100 Hz: median = 0.49, V = 3499.5, p = 2.42e-07; 50 Hz: median = 0.95, V = 5890, p = 1.03e-14; 10 Hz: median = −0.051, V = 2455.5, p = 0.52, Figure 2B-D). These data indicate that extracellular calcium influences short-term synaptic plasticity at the MNTB-MOC synapse.

### Calcium dependence of recovery from short-term plasticity

Following a period of sustained activity that elicits short-term synaptic plasticity, synaptic responses typically recover to baseline levels with the recovery time indicating when synaptic responses will return to their maximal effect. We investigated recovery of MNTB-MOC synapses after synaptic depression following trains of electrical stimulation of MNTB axons of 20 pulses at 100 Hz, as above, and also probed the calcium dependence of recovery in extracellular calcium concentrations of 1.2, and 2.0 mM. The amplitude of PSCs evoked by MNTB stimulation were normalized to the first PSC in the train. The facilitation index of the final PSC in the train showed significant depression, as above (1.2 mM calcium: index = −0.56 ± 0.32, Wilcoxon Rank Sum Test W = 34, Z = 2.17, p = 0.030, n = 8 neurons; 2.0 mM: index = −0.68 ± 0.29, Wilcoxon Rank Sum Test W = 50, Z = 2.24, p = 0.025, n = 10 neurons, Figure 3A, B). Following the last pulse in the train, MNTB axons were again stimulated after a recovery interval ranging from 20 ms to 15 s (Figure 3A, B). The amplitude of PSCs evoked after the recovery interval were normalized to the first PSC in the original 100 Hz train. Evoked PSCs recovered to baseline values after about 1 second, with a faster time constant of recovery in 2.0 mM compared to 1.2 mM calcium (recovery time constant: 1.2 mM Ca^2+^: 470.10 ± 208.47 ms, n = 7-9 cells per interval; 2.0 mM Ca^2+^: 118.13 ± 65.36 ms, n = 5-7 cells per interval; ePSC ∼ α*exp(Time * β) + ϴ, larger magnitude of β indicates steeper decay: 1.2 mM Ca^2+^ β = −2.1 ± 1.6 SE, 2.0 mM Ca^2+^ β = −35.5 ± 28.1 SE, p = 0.0244, Figure 3C, Ci), demonstrating a calcium dependence of recovery from synaptic depression.

**Figure 3.**
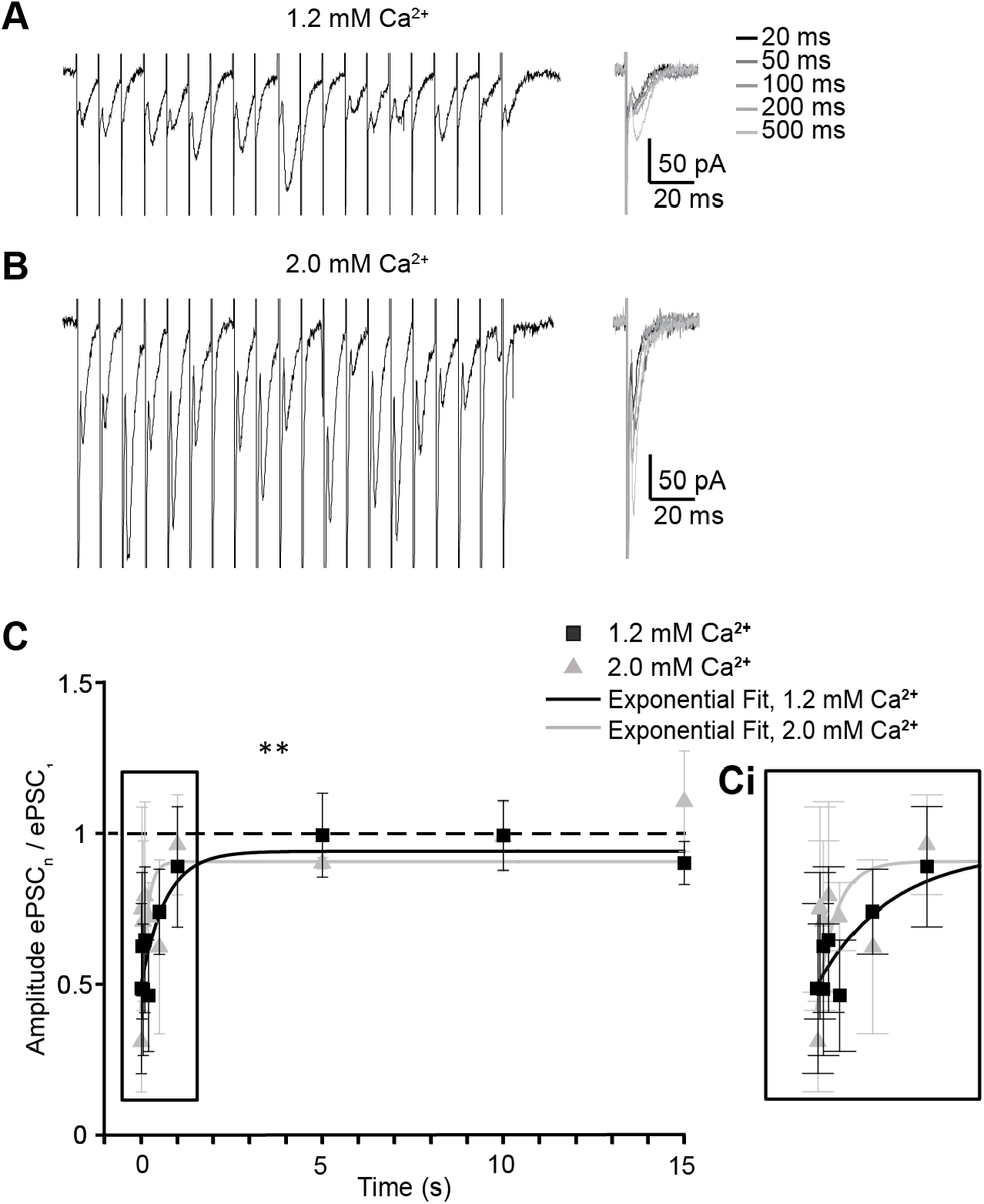
Calcium dependence of recovery of MNTB-MOC synapses from depression. A-B. Example PSCs evoked by 100 Hz stimulation in 1.2 mM calcium, followed by PSC evoked after intervals from 20 ms to 500 ms in 1.2 mM (A) or 2.0 mM (B) extracellular calcium. C. Amplitude of the average PSC evoked after 20 ms to 15 s recovery following 100 Hz trains of stimulation of MNTB axons, normalized to the first ePSC in the train, in 1.2 (black) or 2.0 (grey) mM extracellular calcium. Curves are exponential fits of the data. Ci: zoom of 20 ms to 1 s recovery timepoints from region in ‘C’ indicated by rectangle.

### MNTB-MOC synaptic function following tonic background MNTB activity

MNTB neurons have a high background firing rate in the absence of sound up to 189 Hz, with average rates around 10-20 Hz documented in multiple species (Sommer et al., 1993; Smith et al., 1998; Kopp-Scheinpflug et al., 2003, 2008; Hermann et al., 2007; Kadner and Berrebi, 2008). This high rate of background firing suggests that the MNTB-MOC synapses may be tonically depressed *in vivo*, even in quiet. Therefore, we mimicked the tonic activity level present *in vivo* at the MNTB-MOC synapse to determine if this baseline activity could alter ePSCs in MOC neurons at faster activity rates, such as might occur during the transition from quiet to sound-driven activity. Experiments were performed in 1.2 mM extracellular calcium and at physiological temperature. To mimic tonic firing, we applied “background” stimulation to MNTB axons at 10 Hz, a typical spontaneous rate for MNTB neurons. This firing rate evokes mild synaptic depression at MNTB-MOC synapses, while a high rate of baseline stimulation at 100 Hz evokes strong synaptic depression at the MNTB-MOC synapse, as shown above. After two seconds of mild (10 Hz) or strong (100 Hz) baseline stimulation we then applied a 400 ms stimulation, at 50, 100, 200, or 500 Hz, to mimic enhanced MNTB synaptic activity associated with sound (Figure 4A, B). We compared the peak current amplitude reached in the control condition to the peak current amplitude reached following background stimulation. Mild background stimulation at 10 Hz did not reduce the subsequent responses to faster stimulation, but faster (100 Hz) background stimulation inhibited the response to subsequent stimulation at 200 Hz (Figure 4C-G, Table 1). Thus, if a given MNTB neuron has a typical, mild, rate of background activity, the MNTB synapse will have a significant effect on the MOC neuron at sound onset. However, for the MNTB neurons with a high background activity, further synaptic activity evoked by sound may have a limited effect on the post-synaptic MOC neuron.

**Figure 4.**
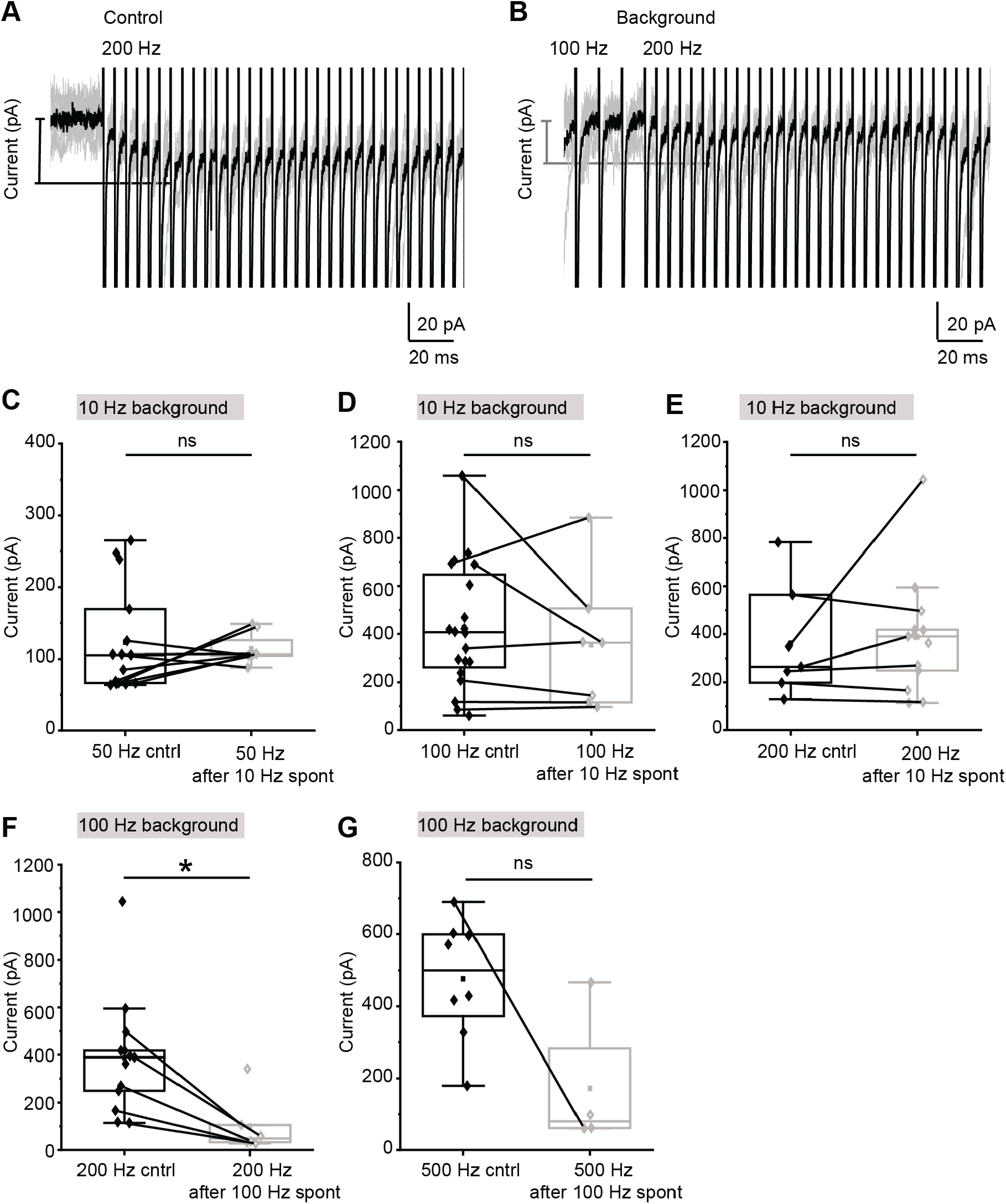
High rates of background MNTB synaptic activity reduce responses to subsequent activity. A. Example voltage clamp trace showing the response of an MOC neuron to 200 Hz stimulation of MNTB axons. Grey: individual traces. Black: average trace. Vertical lines are the stimulation artifact. B. Same neuron as in ‘A’, response to 100 Hz background stimulation of MNTB axons followed by 200 Hz stimulation. C-E. Plots showing the peak current measured as in A-B during trains of MNTB axon stimulation either in control conditions or following 10 Hz background stimulation of MNTB axons. Lines connect datapoints from the same neuron. F-G. Plots showing the peak current measured as in A-B during trains of MNTB axon stimulation either in control conditions or following 100 Hz background stimulation of MNTB axons. Lines connect datapoints from the same neuron.

**Table 1.** Peak current amplitudes at MNTB-MOC synapses in control conditions and following background MNTB stimulation. Fast rates of MNTB axon stimulation were applied at the rates indicated (“sound-like” stimulation rate), either in control conditions or following two seconds of mild (10 Hz) or strong (100 Hz) background stimulation. Data are presented as median ± MAD.

| Background stim rate (Hz) | Fast stim rate (Hz) | Peak amplitude control (pA) | Peak amplitude after background stim (pA) | p-value | n control / background stim |
| --- | --- | --- | --- | --- | --- |
| 10 | 50 | 276.9 +/- 102.7 | 227.2 +/- 172.25 | 0.78 | 15/8 |
| 10 | 100 | 407.05 +/- 182.55 | 363.4 +/- 218.6 | 0.52 | 20 / 7 |
| 10 | 200 | 390 +/- 119.7 | 264.5 +/-83.1 | 0.69 | 13/7 |
| 100 | 200 | 160.74 +/- 119.7 | 81.91 +/- 17.9 | 0.0033 | 13/6 |
| 100 | 500 | 500.35 +/- 99.05 | 80.15 +/- 18.9 | 0.0508 | 8/4 |

### Rapid, sustained activity at MNTB-MOC synapses

Sound-driven responses of MNTB neurons *in vivo* can be up to 500-700 Hz (Kopp-Scheinpflug et al., 2008). Because MNTB-MOC synapses depress during repetitive stimulation, we asked whether these synapses can maintain activity even when rapid stimulation rates are applied for sustained durations. For this work we applied trains of electrical stimulation to MNTB axons at rates of 200 and 500 Hz for a duration of 1 s. During this sustained stimulation we found that evoked responses summated and became indistinct over the duration of the trains to reach a peak amplitude that incorporated both summation and short-term plasticity (Figure 5A, B). The peak amplitude was not different between 200 and 500 Hz stimulation rates (peak amplitude: 200 Hz: 395.90 ± 158.5 pA, n = 5 neurons; 500 Hz: 519.25 ± 225.3 pA, n = 6 neurons, Mann-Whitney U test: U = 13, Z = −0.27, p = 0.78, Figure 5C). Although there was no change in peak amplitude, there was a trend towards a faster latency from stimulation onset to peak in MNTB-MOC synapses stimulated at 500 compared to 200 Hz (latency to peak: 200 Hz: 45.36 ± 11.09 ms, n = 5 neurons; 500 Hz: 26.44 ± 8.46 ms, n = 6 neurons, Mann-Whitney U test: U = 26, Z = 1.92, p = 0.055, Figure 5D). Following this peak in activity, the total current had an exponential decay with a time constant that was not different between stimulation rates (200 Hz exp decay: 715.92 ± 106.31 ms, n = 5 neurons, 500 Hz decay: 447.23 ± 277.47 ms, n = 6 neurons, Mann-Whitney U test: U = 15, Z = 0.42, p = 0.68, Figure 5E) and also decayed to a plateau current that was not different between stimulation rates (plateau current: 200 Hz: 92.4 ± 35.2 pA, n = 5 neurons, 500 Hz plateau: 144.6 ± 93.8 pA, n = 6 neurons, Mann-Whitney U test: U = 10, Z = −0.82, p = 0.41). Thus, during sustained MNTB-MOC synapse activity such as may occur during a sustained sound, the evoked responses build to a peak followed by decay but are able to maintain a sustained inhibitory current.

**Figure 5.**
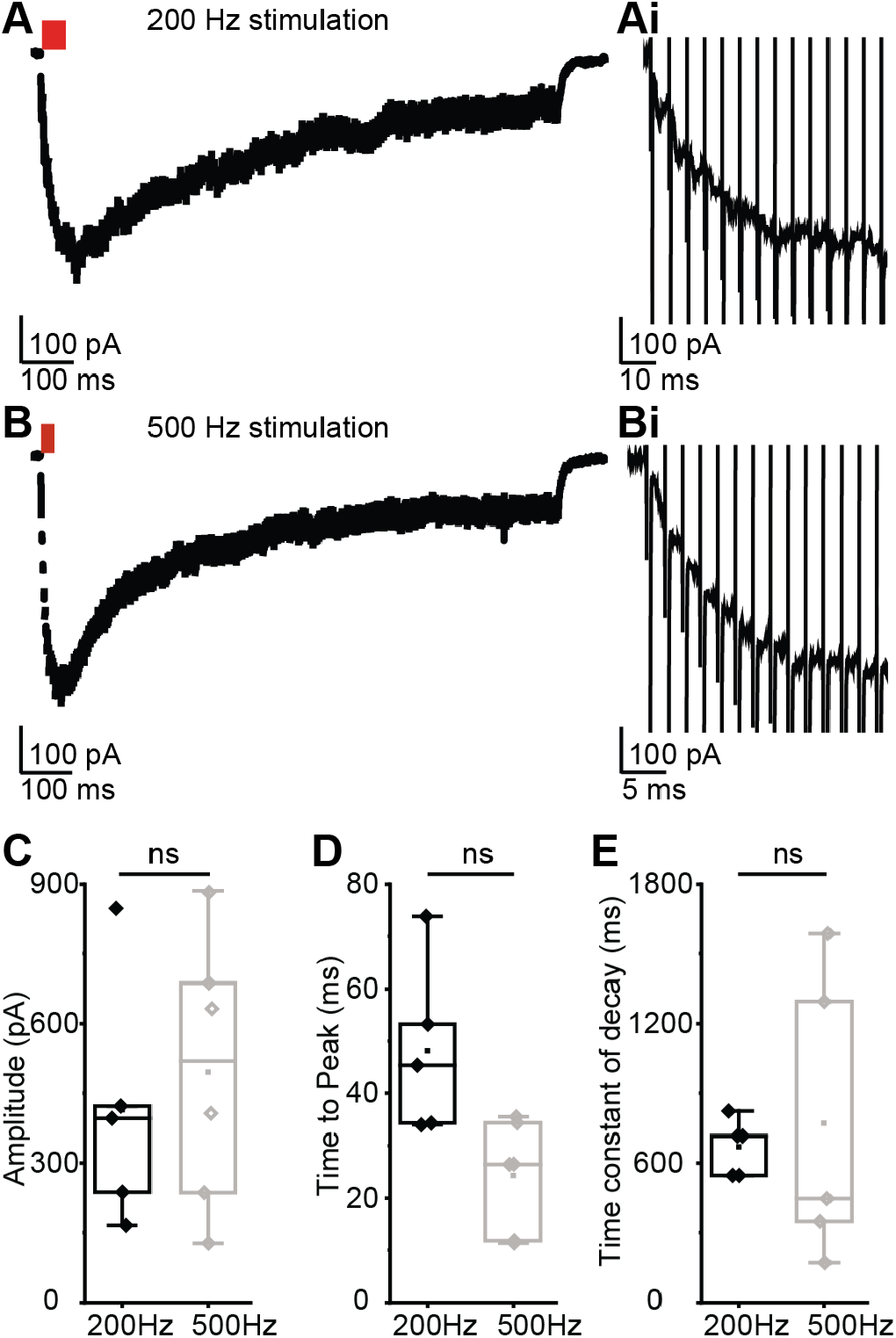
High frequency activation of MNTB-MOC axons evokes strong synaptic depression followed by tonic activity. A. Example trace showing MOC neuron response to 200 Hz stimulation of MNTB axons. Stimulus artifacts were digitally removed. Red indicates region of the trace used in zoom in Ai. B. Example trace showing response to 500 Hz stimulation of MNTB axons. Stimulus artifacts were digitally removed. Red indicates region of trace used in zoom in Bi. C. Plot of peak amplitude of summated ePSCs evoked by stimulation of MNTB axons at 200 or 500 Hz. D. Plot of time to peak of summated ePSCs evoked by stimulation of MNTB axons at 200 or 500 Hz. E. Time constant of decay of summated ePSCs evoked by stimulation of MNTB axons at 200 or 500 Hz.

## Discussion

Functional evidence describing the complexity of synaptic inputs to MOC neurons corroborates existing histological evidence that multiple classes of neurons form synapses onto olivocochlear efferent neurons, indirectly influencing cochlear function. Recent work demonstrates that MOC neurons are excited by both ascending sound-driven inputs from CN T-stellate cells, and descending excitation from the IC (Hockley et al., 2021; Romero and Trussell, 2021). MOC neurons are also inhibited by ascending sound-driven pathways via MNTB neurons (Torres Cadenas et al., 2020). There are likely additional sources of excitation, inhibition, or modulation from auditory and other brain regions as indicated by histological studies (Faye-Lund, 1986; Caicedo and Herbert, 1993; Thompson and Thompson, 1993; Vetter et al., 1993; Mulders and Robertson, 2000, 2002; Groff and Liberman, 2003; Horvath et al., 2003; Ota et al., 2004; Gómez-Nieto et al., 2008b; Brown et al., 2013; Suthakar and Ryugo, 2016). The integration of these synaptic inputs to govern MOC activity depend on the relative strength and post-synaptic effect of inputs, and their plasticity during changing hearing conditions. Here we further characterize the short-term plasticity of inhibitory inputs to MOC neurons from neurons of the MNTB, in mouse. MNTB-MOC synaptic inputs depress over a range of physiological calcium and stimulation rate conditions, yet maintain sustained synaptic function. This holds even during high rates of activity and following background spontaneous rates of activity.

### Temperature and calcium effects on MNTB-MOC plasticity

Increasing the temperature of tissue speeds the kinetics of biological processes, including those regulating vesicular neurotransmitter release (Sabatini and Regehr, 1996). Consistent with this idea, raising the temperature of a brain slice from RT to PhT during patch-clamp recordings of MNTB-MOC synaptic responses both decreased the latency and sped up the kinetics of evoked PSCs. Using PhT during recording has also been shown to decrease the latency and speed of evoked PSCs in neurons from other auditory and non-auditory brain areas (Zhang and Trussell, 1994; Taschenberger and von Gersdorff, 2000; Kushmerick et al., 2006). However, we saw no change in the amplitude of single evoked PSCs in PhT compared to RT. In other cell types an increase in evoked PSC amplitudes with increased temperature has been observed (Zhang and Trussell, 1994; Hardingham and Larkman, 1998; Taschenberger and von Gersdorff, 2000; Kushmerick et al., 2006), sometimes also associated with an increase in the mini-PSC (mPSC) amplitude (Kushmerick et al., 2006; Van Hook, 2020). In other systems that also have an increase in evoked PSC amplitude with increased temperature, this effect was not associated with an increase in mPSC amplitude (Hardingham and Larkman, 1998). It is unclear why we do not see an increased PSC amplitude with temperature. This may simply reflect a lack of change in quantal size with temperature combined with fewer release sites from MNTB onto MOC neurons, especially compared to other studies discussed here that examined synapses with many release sites.

Changes in short-term synaptic plasticity with temperature have been observed at other central synapses. While many mechanisms affect both pre- and post-synaptic plasticity, net plasticity can be in part attributed to a balance between release probability and vesicle recruitment. Release probability has been observed to increase with temperature at many (Hardingham and Larkman, 1998; Pyott and Rosenmund, 2002; Volgushev et al., 2004) but not all (Allen and Stevens, 1994) synapses and is typically associated with increased synaptic depression. Increased vesicle recruitment occurs with increased temperature (Pyott and Rosenmund, 2002; Kushmerick et al., 2006), and is associated with decreased synaptic depression. However, at many synapses the net effect is a reduction in short-term synaptic depression at PhT relative to RT (Brenowitz et al., 1998; Hardingham and Larkman, 1998; Taschenberger and von Gersdorff, 2000; Pyott and Rosenmund, 2002; Volgushev et al., 2004; Kushmerick et al., 2006). Consistent with these studies, we find a decrease in short-term depression at MNTB-MOC synapses when the temperature of the slice is raised to PhT, extending this principal to inhibitory synapses.

In addition to temperature effects, short-term synaptic plasticity involves interplay between many mechanisms, including pre-synaptic calcium dynamics, vesicle and release site properties, neurotransmitter diffusion and clearance from the synaptic cleft, and post-synaptic receptor dynamics, among other mechanisms. Various mechanisms for the strong influence of extracellular calcium on pre-synaptic mechanisms of plasticity have been proposed, including vesicle pool replenishment or depletion, changes in vesicle release probability, and effects on pre-synaptic calcium channels (Sakaba et al., 2002; Zucker and Regehr, 2002; Catterall and Few, 2008; Neher and Sakaba, 2008; Fioravante and Regehr, 2011; Mochida, 2011; reviewed in Regehr, 2012). In our previous work there was no difference between short term depression at MNTB-MOC synapses in 2.0 vs 1.2 mM calcium when measured with paired pulse (PP) stimulation (Torres Cadenas et al 2020), suggesting that synaptic plasticity is independent of extracellular calcium at this synapse. While unusual, this has been observed at other auditory synapses in the CN that were tested with high vs physiological (lower) divalent cations in aCSF (Xie and Manis, 2017). However, extending the duration of synaptic activity to trains of stimulation of MNTB-MOC synapses demonstrated less synaptic depression in 1.2 compared to 2.0 mM calcium. With a further decrease in the calcium concentration to 0.8 mM, MNTB-MOC synapses facilitated, confirming the calcium dependence of plasticity at the MNTB-MOC synapse via an unknown mechanism at the MNTB axon terminal.

The recovery of a synapse from depression is also calcium-dependent, including dependence on mechanisms such as vesicle pool refilling (Dittman and Regehr, 1998; Neher and Sakaba, 2008). Indeed, MNTB-MOC synapses recovered to baseline levels faster in higher calcium conditions. Interestingly, the time constant of recovery from depression of inhibitory synapses to MOC neurons was faster than that of excitatory synapses to MOC neurons from either T-stellate or inferior colliculus neurons (Romero and Trussell, 2021). This suggests that if excitatory and inhibitory inputs to MOC neurons experience similar activity rates, inhibitory synapses will recover from depression faster than excitatory synapses.

### Effect of spontaneous background activity on MNTB-MOC synaptic inhibition

The GBC-MNTB pathway that inhibits MOC neurons is remarkably fast and powerful, with high-fidelity synaptic transfer in response to stimulation up to ∼800 Hz at the calyx of Held synapse *in vitro* (Spirou et al., 1990; Wu et al., 1993; Taschenberger and von Gersdorff, 2000; Futai et al., 2001; Kopp-Scheinpflug et al., 2003; Joshi et al., 2004). *In vivo*, the background spontaneous rate of MNTB activity in the absence of sound averages about 10-20 Hz and can range up to 190 Hz (Sommer et al., 1993; Smith et al., 1998; Kopp-Scheinpflug et al., 2003, 2008; Kadner et al., 2006; Hermann et al., 2007). In response to sound stimulation, MNTB neurons have a primary-like with notch firing pattern (Guinan and Li, 1990; Smith et al., 1998; Paolini et al., 2001; Kopp-Scheinpflug et al., 2008; Tolnai et al., 2009) and spike at rates of up to hundreds of Hz *in vivo* (Kopp-Scheinpflug et al., 2008). Work from other labs has demonstrated that the calyx of Held synapses onto MNTB neurons have smaller evoked responses and more action potential failures following spontaneous background activity, but exhibit less synaptic depression during fast trains (Hermann et al., 2007). In addition, there is a transient synaptic facilitation following background stimulation (Müller et al., 2010). The *in vivo* activity patterns of MNTB-MOC synapses are unknown, but we assumed that, similar to the MNTB soma, they may have some baseline spontaneous activity that evokes tonic depression, followed by faster rates of activity in response to sound. MNTB inhibition of MOC neurons was not reduced by a low, 10 Hz background synaptic stimulation rate, but was reduced by a prior high, 100 Hz synaptic stimulation rate. Therefore, most MNTB neurons with low background spontaneous rates will likely inhibit MOC neurons upon sound onset. In contrast, MNTB neurons with high background activity levels may instead exert a tonic inhibition of MOC neurons that is insensitive to sound onset.

### Effect of inhibition on MOC activity

MOC neurons in guinea pig and cat rarely fire action potentials in quiet *in vivo*. In response to sound they have low threshold responses that occur with variable latencies (Robertson, 1984; Robertson and Gummer, 1985; Liberman and Brown, 1986; Liberman, 1988). In contrast, in young mice (P14-P23), post-hearing onset MOC neurons sometimes have spontaneous action potentials *in vitro* (Torres Cadenas et al., 2020). MOC neurons recorded from brain slices of mice older than those used in the present study (P30-P48) lack spontaneous action potentials (Romero and Trussell, 2021). Activity from inhibitory MNTB inputs is likely silent in these more mature experiments because spontaneous activity from MNTB neurons is largely absent in *in vitro* brain slices (Hermann et al., 2007). This suggests that intrinsic electrical properties, not inhibition, are likely responsible for the low spontaneous spike rate of mature MOC neurons *in vivo*, although evoked inhibition from the MNTB persists in the mature MOC neurons at the same levels as in younger animals (Torres Cadenas et al., 2020). Instead, tonic inhibition from spontaneously active MNTB neurons *in vivo* likely provides electrical shunting that inhibits action potential generation in response to sound-driven inputs. In addition, synaptic inhibition from the MNTB may increase or add variability to the latency of sound-driven action potentials in MOC neurons. Thus, inhibitory synapses from the MNTB to the MOC are likely characterized by tonic activity that under most conditions of low background activity would continuously inhibit MOC function in quiet, and robustly inhibit MOC neuron function at sound onset. These functions combine to delay MOC suppression of cochlear activity during rapidly modulating sound.

## Acknowledgements

This research was supported by the Intramural Research Program of the NIH, NIDCD, Z01 DC000091 (CJCW).

## Notes

<u>Conflicts of Interest</u>: The authors declare no competing financial interests

### Competing Interest Statement

The authors have declared no competing interest.

